# Lumbar intrathecal catheterization in rats targeting the cerebral cortex: a drug delivery method and validation

**DOI:** 10.64898/2026.06.01.727192

**Authors:** Omar S. Elwardany, Abdiel Badillo-Martinez, Bassel Awada, John L. Bixby, Vance P. Lemmon, Hassan Al-Ali

**Affiliations:** Miami Project to Cure Paralysis, University of Miami, Miami, FL, USA; Department of Biochemistry and Molecular Biology, University of Miami, Miami, FL, USA; Department of Neurological Surgery, University of Miami, Miami, FL, USA; Frost Institute for Data Science and Computing, University of Miami, Miami, FL, USA; Sylvester Cancer Comprehensive Center, University of Miami, Miami, FL, USA; Department of Molecular and Cellular Pharmacology, University of Miami, Miami, FL, USA; Department of Medicine, University of Miami, Miami, FL, USA; Peggy and Harold Katz Family Drug Discovery Center, University of Miami, Miami, FL, USA

**Author notes:** Correspondence: Vance P. Lemmon,; Hassan Al-Ali.

**Keywords:** intrathecal delivery, target engagement, drug delivery, kinase inhibitor, blood-brain barrier

## Abstract

Intrathecal (IT) drug delivery is a critical technique for bypassing the blood-brain and blood-spinal cord barriers in preclinical CNS research. However, conventional rat catheterization methods suffer from high rates of neurologic complications, poor reliability, and unverified dosing due to epidural reflux and inconsistent supraspinal distribution. Our objective was to develop and validate an improved method for lumbar IT catheterization in rats that ensures distribution to the brain and to confirm supraspinal pharmacodynamic target engagement. We describe a refined microsurgical technique using dural puncture under direct visual control at the L6-S1 interlaminar space, a site chosen for its anatomical safety margin. The method uses a small-bore (0.33 mm OD) polyurethane (PU) catheter to minimize durotomy size, air-bubble tracking to compensate for catheter dead volume, and epidural sealing with Surgifoam^®^ to minimize reflux. Visualization using Evans Blue dye confirmed complete neuraxial distribution from a single 30 µL lumbar bolus injection, with dye reaching the ventral/dorsal brain cisterns. Pharmacodynamic validation of cortical exposure was achieved using an S6 kinase 1 (S6K1) inhibitor. Lumbar IT administration over a period of 6 hours via a pump resulted in significant supraspinal S6K1 engagement, demonstrated by a reproducible reduction in S6 phosphorylation in the cerebral cortex.

**Highlights:**

- Method for lumbar intrathecal catheterization in rats under direct visual control, using basic surgical tools
- CNS distribution validated by Evans Blue dye reaching ventral and dorsal brain
- Pharmacodynamic confirmation of supraspinal target engagement following lumbar intrathecal delivery of a small molecule kinase inhibitor
- Serves as a faithful preclinical model for therapeutics intended for clinical intrathecal administration
- Provides a screening route for early-stage compounds not yet optimized for CNS penetrance, supporting efficacy testing prior to medicinal chemistry investment

## Introduction

Treatment of CNS disorders is constrained by the BBB and the blood-spinal cord barrier, which together restrict over 98% of potential neurotherapeutics [1-3]. Achieving therapeutic CNS concentrations via systemic administration, therefore, often requires doses with prohibitive toxicity. This impasse has driven direct-to-CNS strategies, with IT administration into the CSF emerging as a clinically translatable route bypassing these barriers [4]. Utilizing this route in preclinical research provides a versatile evaluative framework. For therapeutics explicitly intended for clinical intrathecal administration, the procedure provides a faithful model of the clinical delivery route. For early-stage compounds ultimately intended for systemic administration but not yet optimized for CNS penetrance, this approach allows them to be tested for efficacy, justifying further development.

Rats are a preclinical model for this delivery route [5-7], which exploits CSF flow for drug distribution, including to supraspinal regions [8]. In preclinical settings, rodent IT catheterization also provides a means to access CSF for repeated drug delivery and/or sampling. The technique, pioneered by Yaksh and Rudy in 1976, established a foundation for spinal cord pharmacology and neuroscience studies in rats using lumbar catheterization [5].

Several refinements have been proposed. A lumbar approach with PE10 catheters and guide cannulas is reliable in rats [9]. A microvalve catheter improves distribution along the spinal axis, avoids proximal pooling, reduces obstruction, and enables transfection of cervical-to-lumbar dorsal root ganglia after viral infusion [10]. A “needle-through-needle” design improves practicality and stability, lowering mortality with performance comparable to laminectomy and fewer complications [11]. Despite these modifications, existing techniques still carry significant risks of spinal cord injury, subarachnoid hemorrhage, postoperative sensory/motor deficits, mortality, and procedural unreliability [7, 11, 12].

Our method, utilizing basic surgical tools (for a complete list, see Online Resource 1), a small diameter catheter inserted at the L6/S1 inter-laminar space under visual control, along with adopting “CSF backflow” as the sure sign of successful catheterization, has been validated with two infusates. Evans Blue dye delivered as a bolus reached the ventral and dorsal brain CSF spaces, and an S6 Kinase 1 (S6K1) inhibitor, compound 8851, engaged its cortical target after continuous 6-hour intrathecal infusion via programmable pump. This method addresses an unmet need in preclinical CNS delivery.

## Methods

### Animals & Ethics

In vivo procedures followed protocol 21-138 approved by the University of Miami IACUC. The University’s animal research program is AAALAC International-accredited.

Adult male Sprague-Dawley (dye injection) or Long Evans (target engagement) rats (300-400 g) were housed 2 per cage in a temperature-controlled (20 °C ± 2 °C) and light-controlled (12-h light/dark cycle) room, with ad libitum access to food and water.

### Materials and Reagents

Compound 8851, an S6K1 inhibitor developed in-house (US Patent Application US 2025/0145586 A1), was selected for its demonstrated brain exposure after intrathecal delivery in rats and is available to qualified researchers under MTA. Stock solution of compound 8851 in dimethyl sulfoxide (DMSO) was diluted with D5W (5% dextrose in water; Gibco, cat#15023-021), then Kolliphor HS 15 (BASF, cat#50259817), preheated to liquefy from its solid form, was added to achieve a final concentration of 30 mM compound 8851 in 10% DMSO, 89% D5W, and 1% Kolliphor HS 15. Vehicle controls were identical without active compound. All formulations were sterile-filtered (0.2 µm nylon; ThermoFisher, cat# 726-2520). Evans Blue (Sigma-Aldrich, E2129-10G, Lot#00801KHV) was made to 1% in normal saline and filter sterilized. 8% SDS Buffer: tris-HCl (pH 6.5; 0.2 M), glycerol (Sigma-Aldrich, cat# G15516; 32% v/v), SDS (Sigma-Aldrich, cat# L3771; 8% w/v), bromophenol blue (Sigma-Aldrich, cat# B0126; 0.04% w/v).

### Equipment

Rat Intrathecal Catheter: Polyethylene (PE-10) tubes (ID 0.28 mm; OD 0.61 mm) are most commonly used for rat IT catheterization [9, 11]. We initially used PE-10 but identified drawbacks. First, sterile PE-10 was unavailable from vendors. Second, the large OD (0.61 mm) is a disadvantage in rodents. Third, we and others [9] used 0.009” OD PTFE guide wires for PE-10, which were sold only non-sterile. We therefore switched to Polyurethane (PU) Rat Short Intrathecal Catheters (OD 0.33 mm; Alzet No. 0007741). Despite being more technically demanding to use, such catheters come sterile and singly packed along with a preloaded Teflon-coated stainless-steel stylet (OD 0.005”) for easier placement.

### Programable Micro Pumps for infusion

To deliver test compound over extended periods or in multiple doses, we used iPrecio programmable micro infusion pumps (SMP-310R, Alzet). These pumps are sterile, individually packed, and can be programmed to infuse a defined volume over a defined period. The pump was connected to the lumbar IT catheter after confirming catheter position and stabilization (see Technical Observations). Once connected, the pump was placed in a subcutaneous flank pocket prepared during the initial skin incision, adjacent to the lumbar incision.

### Hamilton syringe for direct bolus injection

Initial bolus delivery used a 50 µL glass syringe (Hamilton, 705RN 40585), connected to the IT catheter via a transitional tube due to needle/catheter mismatch. Recently, we started using a custom syringe (Hamilton, 710 SN/22G/1IN/45GED 100 µL, 80608). This syringe/needle provided an easy and secure connection of the needle directly to the distal end of the IT catheter (see technical notes below). Injection using the syringe was managed with a QSI Dual Microliter Syringe Pump, Tabletop System (by Stoelting) to infuse 20-40 μL at a rate of 5-8 μL/min (see technical notes below).

### Intrathecal Compound Delivery for CNS Targeting

To validate direct-to-CNS delivery, we administered compound 8851 intrathecally. This approach was chosen to bypass the BBB and ensure access to neural tissues, facilitating drug distribution to neuronal targets. Importantly, IT drug administration minimizes the need to traverse additional biological barriers, enhancing bioavailability at the intended sites of action. For optimal safety and efficacy, we selected the most caudal interlaminar space (L6-S1) for injection. This represents the furthest accessible point from the conus medullaris where a dural puncture can be performed safely. To further mitigate potential neurotoxicity and cord damage, we maintained the catheter length within the thecal sac at 1–1.5 cm. This short catheter length, combined with its L6-S1 placement, minimizes exposure of the caudal spinal cord to the highest concentration of injected compound while allowing adequate mixing with CSF, minimizing direct neurotoxicity. Additionally, positioning the catheter tip distal to the spinal cord minimizes the likelihood of inadvertent cord injury during catheter placement. Our procedure thus ensures effective compound distribution to the relevant CNS regions, including, in cases of spinal cord injury, the spinal cord injury site as well as the cortex and other brain structures, without compromising mechanical stability.

### Anesthetic protocol

Both inhalational (isoflurane) and parenteral anesthetics (ketamine/xylazine given as intraperitoneal (IP) injections) were used. Extended-release buprenorphine (preoperative SC, single dose) was given for survival surgeries (iPrecio pump implants).

### Intrathecal Catheterization with Pump Implantation or Direct Syringe Injection

After anesthesia, the lumbosacral back was shaved and wiped with Nolvasan. The rat was placed in a prone position on a warm pad controlled via a rectal temperature probe. Using a surgical microscope (DF Vasconcellos), a midline skin incision of 1.5 – 2 cm over the lumbosacral area was made. After clearing the subcutaneous tissue, the sheath of paravertebral muscles was sharply incised bilaterally with a micro-scissor to expose the spinous processes of L6 and S1 vertebrae. The L6-S1 supraspinous and interspinous ligaments were cut to provide midline access to the interlaminar space. The nearby bilateral muscles were retracted laterally until the dura at the L6/S1 interlaminar space was clearly visualized. An intrathecal catheter was inserted under visual control, piercing the dorsal dura at the L6/S1 level (either utilizing the tip of the preloaded guidewire or after creating a dural puncture using the tip of a 23G needle). After confirming catheter position (see Technical Observations), the distal end was connected to an iPrecio SMP-310R pump or to a 22G Hamilton syringe needle (Figure 1). The pump was implanted in a subcutaneous flank pocket. For bolus injection, the catheter was held in place 1-2 minutes after delivery before withdrawal. After muscle sheath closure with 2-3 absorbable sutures, the skin incision was closed using absorbable/non-absorbable sutures or surgical skin clips.

**Figure 1:**
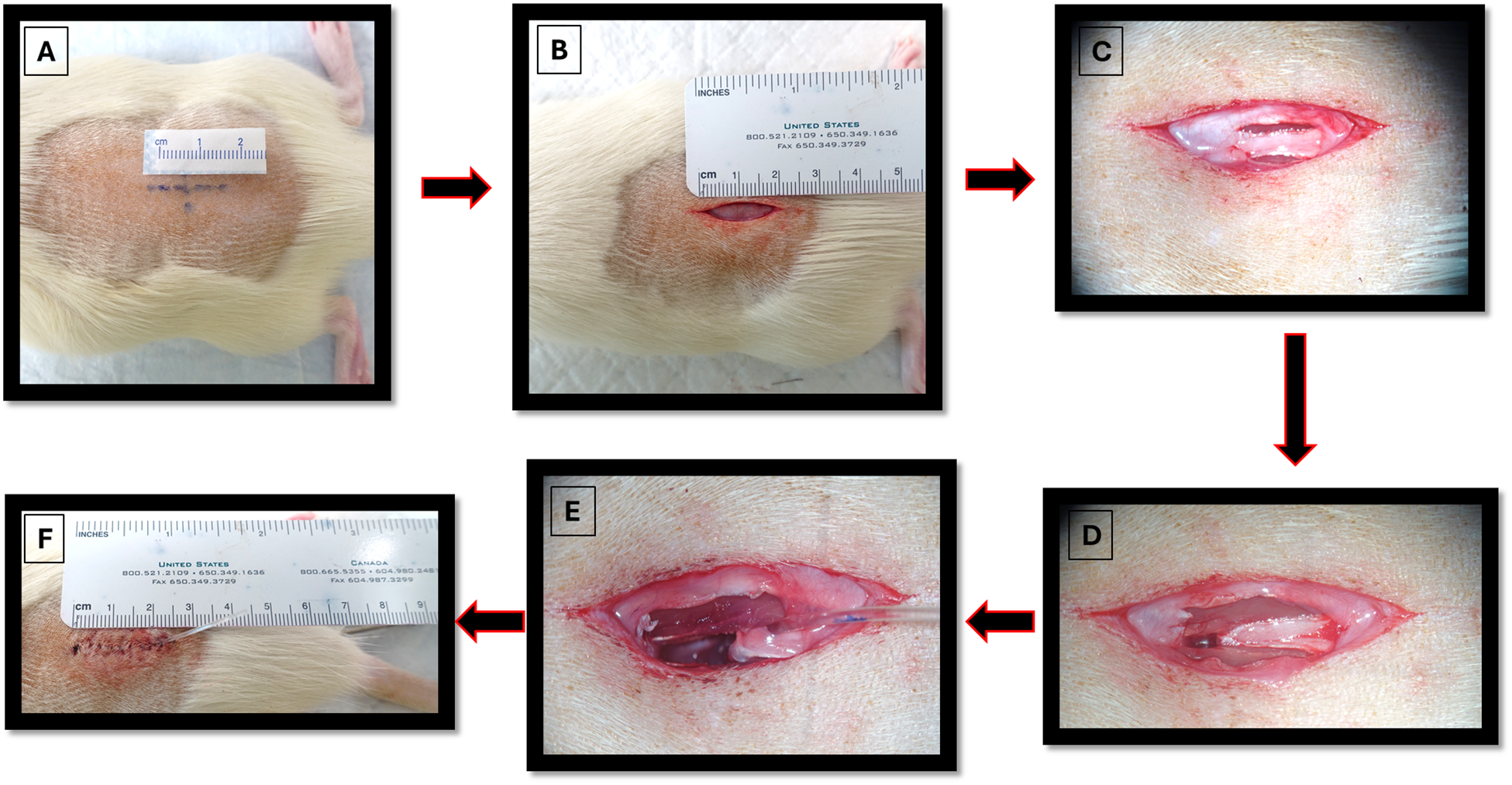
A: Skin marking for the incision centered on L6/S1 inter-spinous level. B: Mid-line skin incision (2 cm). C: Bilateral cut of the paravertebral muscle sheath. D: cutting L6/S1 supra-spinous and inter-spinous ligaments. E: The catheter piercing the dura at L6/S1 inter-laminar space. F: After multi-layer closure.

One surgeon performed all preoperative preparations and surgery (pump programming, loading, activation; IT catheterization; pump implantation), assisted by 1-2 team members for animal preparation, note-taking, and post-surgical observation. A second team of 2-3 people sacrificed the animals and harvested the tissues for Western blots.

### Target Engagement Experiment

Male Long Evans rats underwent pump implantation on day 1 and were allowed to recover overnight. The pump was preloaded with a 30 mM solution of compound 8851 and programmed to start infusing the next morning at a rate of 3 μL/hr (total ∼ 1 mg/kg). Six hours after infusion start, the animals were sacrificed and the CNS was rapidly dissected. The brain was rinsed with ice cold PBS and placed on an ice-cold dissection surface. From one hemisphere, a full-thickness cortical block (∼100 mg wet weight) was excised with a scalpel, extending from midline to lateral cortical edge and centered anteroposteriorly between caudal olfactory bulb and rostral brainstem (∼bregma 0 to -3 mm). The dissection included the full cortical mantle (layers I–VI) without underlying white matter or subcortical tissue. The cortical block was homogenized in 250 μL of hot 8% SDS Sample Buffer containing complete™, Mini, EDTA-free Protease Inhibitor Cocktail (PIC) (Roche, 118366170001). After homogenization, another 750 μL of SDS Sample Buffer with PIC was added, and the sample was left in a 100 °C heat block for 10 min. The samples were centrifuged at 20,800×g for 5 min in an Eppendorf Centrifuge 5417R, and the supernatant was stored at -80 °C until Western blot was performed.

### Western Blots

NuPAGE Bis-Tris Mini Protein Gels (4-12%) (ThermoFisher #NP0321BOX) were used along with NuPAGE MOPS SDS running buffer (ThermoFisher #NP0001) per manufacturer. Samples were heated at 100 °C for 2 min, centrifuged at 20,800×g for 2 min, mixed with beta-mercaptoethanol (1:25), and 20 µL was loaded per lane. After the gels were run, the proteins were transferred to nitrocellulose (Millipore, cat# IPVH00010) using NuPAGE Transfer Buffer (ThermoFisher #NP00061) with 25% methanol. The membrane was blocked in Licor Blocking Buffer (Fisher Scientific, cat# NC1660550) for ≥1 h at room temperature (RT), then incubated overnight at 4 °C with primary antibodies in blocking buffer. The membrane was washed 3×5 min in Tris Buffered Saline with 0.1% Tween 20 (TBST) and incubated in secondary antibodies in blocking buffer for 1 hour at RT on a shaker. After 5×5 min TBST washes, the nitrocellulose was imaged on an Azure c600 imager with AzureSpot Pro 1D v1.4 (rolling-ball background correction, automatic band detection, fixed lane widths). The Western blot signals of compound 8851 treated samples were normalized to the control samples in the same nitrocellulose blot.

### Antibodies

1) Primary antibodies: Mouse mAb anti-S6 Ribosomal Protein (Cell Signaling Technology cat# 2317, RRID: AB_2238583) (1:500). Rabbit mAb anti-Phospho-S6 Ribosomal Protein (Ser240/244) (Cell Signaling Technology cat# 5364, RRID: AB_10694233) (1:1,000).

2) Secondary antibodies: 680 RD Donkey anti-Rabbit (LICORbio cat #926-68073, RRID: AB_10954442) (1:10,000). 800 CW Donkey anti-Mouse (LICORbio cat#926-32212, RRID: AB_2716622) (1:5,000).

### Evans Blue Dye Experiment

Injection of Evans Blue (1% in normal saline) was used to validate the ability of our modified method to target CNS tissues generally. Nine-week-old male Sprague-Dawley rats (300-400 g) were used. After catheterization and confirmation of catheter position (CSF seen slowly seeping from the catheter end), the Evans Blue syringe was connected and 40 µL (30 µL + 10 µL dead space) was injected at 6 µL/min using a motorized digital micro-injector. Ninety minutes after injection, rats were euthanized by CO2 chamber followed by thoracotomy and right atrium opening; brain and spinal cord were immediately dissected. Low magnification photos were taken of the exposed spinal cord (after full laminectomy from C1 to L6) and both dorsal and ventral brain (after cranial vault removal)

### Blinding

In the target engagement experiment, the individual filling the syringes for injection and the surgeon were blinded to the identity of the solutions (compound and vehicle), as was the team performing the dissection and western blotting.

## Results

### Our process eliminates the need for a pad beneath the rat during catheterization

Pelvic padding under the rat’s ventral aspect during IT catheterization is common practice [9, 11]. It induces lumbosacral flexion that eases rostral catheter navigation. In our study, gentle upward traction on the L6 spinous process produced sufficient lumbar curvature for smooth rostral navigation. Omitting the pad maintains normal intra-abdominal and pelvic pressure, avoiding the paraspinal venous congestion that bloodies the surgical field, obscures the catheter’s dural entry point, and prolongs hemostasis.

### A sure sign of proper access to the intrathecal compartment (subarachnoid space)

Tail flicks or hind limb twitches during catheter insertion are widely accepted indicators of successful entry into the thecal sac [9, 11]. With our distal site (L6-S1) and short catheter length, these signs were often absent despite clear CSF backflow. While we observed these signs, we considered the immediate appearance of CSF backflow upon removal of the guide wire as the definitive confirmation of successful catheter placement. When backflow was not immediately evident (typically after multiple dural punctures, likely from transient CSF leakage), gentle abdominal pressure usually drew CSF into the catheter. A progressively ascending CSF column confirmed correct intrathecal placement. Absence of backflow indicated a false passage, requiring withdrawal and reinsertion; catheter replacement was needed only if the catheter suffered kinking or structural damage. Occasionally, blood from the epidural venous plexus entered the catheter tip and obstructed backflow; we then withdrew the catheter, removed the stylet, flushed with normal saline, and ensured field hemostasis before re-catheterization. Bloody CSF was sometimes observed in the catheter; as long as flow rate was normal, it was deemed clinically insignificant, likely from minor dural capillary disruption, and did not affect catheter function or procedural integrity.

### A reliable way to ensure the exact desired intrathecal catheter length

Some investigators mark the catheter’s distal limit with a dye-based marker to visualize it during advancement in the thecal sac [9]. No safety data exist for use of dye-based markers in the CSF or proximal to the CNS. We instead exploited the Alzet catheter’s built-in protective tubing junction (designed to minimize kinking at the IT exit point); the catheter tip was trimmed to leave only 1.5 cm distal to the junction (the optimal intrathecal length). By doing this, we could determine the exact intrathecal length of the catheter during initial placement. In addition, because of the larger OD of the tubing junction, we could prevent any later inadvertent advancement of the catheter into the thecal sac.

### Direct visualization of the dura provides accurate dural entrance

Many rodent IT studies insert the catheter through the dura using surface landmarks (as in the clinic) without exposing the vertebra; this less-invasive approach carries higher failure risk from inadequate placement. In our study, we sought to adopt the “under visual control” method of placing the IT catheters. This strategy uses a small window, created under a surgical microscope. The supraspinous and interspinous ligaments between L6 and S1 are snipped, followed by a blunt dissection of the muscles to free the midline, where the dura and the overlying ligamentum flavum can be seen.

### Fixing the IT catheter in place via paravertebral muscle sheath closure

After confirming catheter position, two to three interrupted sutures were inserted to close the paravertebral muscle sheath proximal and distal to the catheter exit. The closed sheath retains the larger OD tubing junction, stabilizing the catheter without circumferential sutures.

### Minimizing efflux of injected compound into the open epidural space

For intraparenchymal injections, it is common practice to leave the needle or catheter in place 1-3 minutes after delivery, allowing the injectate to diffuse and minimizing efflux from the target site, thereby enhancing delivery of the desired volume. Beyond this, we minimized efflux by filling the exposed epidural and interlaminar space at the catheter site with a small piece of Surgifoam^®^ (Ethicon, ref. 1977). As the Surgifoam soaks with blood, it occupies the space and minimizes backflow of injectate or CSF after catheter removal.

### A reliable way to ensure delivery of the desired volume of injected compound (minimizing dead volume)

Direct bolus injection via the IT catheter provides reliable CNS-targeted delivery, and the catheter’s flexibility allows use of a motorized digital injector for precise control. However, an inherent technical challenge arises from the catheter’s dead space (i.e., its internal volume), which must be accounted for when calculating the total injected volume, particularly for small-volume administrations. We addressed this by leaving a small air bubble at the catheter’s distal end where the syringe needle connects; as injection proceeds, the bubble advances proximally toward the thecal sac. Monitoring this advance under a microscope confirms drug entry into the IT space and quantifies the dead volume (∼10 µL in our study), which we added to the syringe-pumped amount. Alternatively, the injector can be set to begin delivery of the target dose only after the bubble reaches the catheter’s proximal end within the thecal sac, compensating for the dead volume.

### Only basic surgical tools are needed

Most prior modifications to IT catheterization use additional tools (syringe needles, guide cannulas, spinal/epidural needles, stylets), particularly with needle-through-needle techniques [9, 11]. Our technique utilizes only the basic surgical tools (scalpel, scissors, toothed and non-toothed forceps, and needle holder) in addition to a 23-gauge needle and surgical thread for closure, and is thus simple to implement.

### Target Engagement by kinase inhibitor demonstrated successful delivery to brain

To confirm that the S6K1 inhibitor (compound 8851) reached its cortical target after IT delivery of 18 µL over hours, 100 mg of cortical tissue was harvested immediately after delivery and assessed for phospho-S6 (pS6) by Western blot. The ratio of pS6/total S6 was compared between animals treated with vehicle and with ∼1 mg/kg of compound 8851. The latter produced ∼25% reduction in phosphorylation of S6 at Ser240/244, a specific and reliable marker of S6K1 activity [13], indicating that compound 8851 reached its target in the cerebral cortex when delivered over 6 hours using an infusion pump (Figure 2).

**Figure 2:**
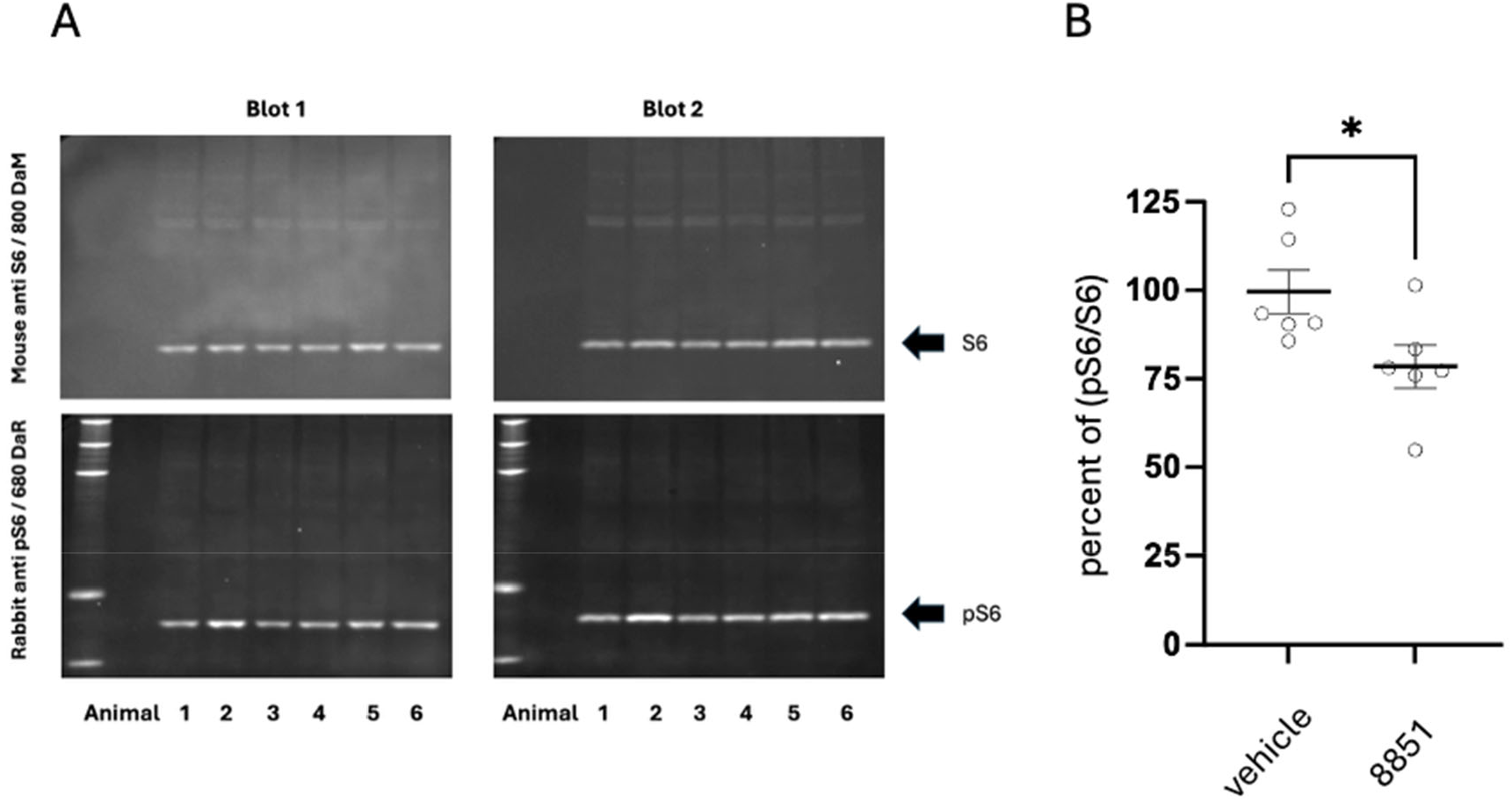
Delivering compound 8851 IT into rats results in detectable reduction of phospho-S6 in the cerebral cortex. compound 8851 was delivered at ∼1 mg/kg as continuous lumbar intrathecal infusion (over 6 hours) in rats. S6 phosphorylation was quantified ∼1.5 hr after dosing using western blotting and expressed as the ratio of pS6 to total S6, normalized to vehicle-treated animals. **A**⍰ Samples from each animal were run on two gels and average areas under the curves calculated by the Azure software were obtained. Two example blots are shown. The top gel shows pS6 and the bottom gel shows S6 protein indicated by the arrows. Animals 1, 3, 5 were treated with compound 8851. Animals 2, 4, 6 were treated with vehicle. **B**⍰ The results from two independent experiments, including a total of 12 animals are shown. Data are expressed as mean ± SEM, N = 6 animals per treatment group. *p = 0.0357 by two-tailed unpaired t-test.

### Evans Blue injection to visualize the extent of distribution of small molecules using lumbar IT injection

Evans Blue (MW 961) tightly binds serum albumin. Because rat CSF albumin (80 µg/mL) is far below serum levels (30-40 mg/mL), Evans Blue is reasonably assumed to act as a small molecule in CSF [14, 15]. Injection via our modified lumbar IT method delivered dye to both ventral and dorsal brain CSF cisterns and the entire spinal cord (Figure 3) within 90 min.

**Figure 3:**
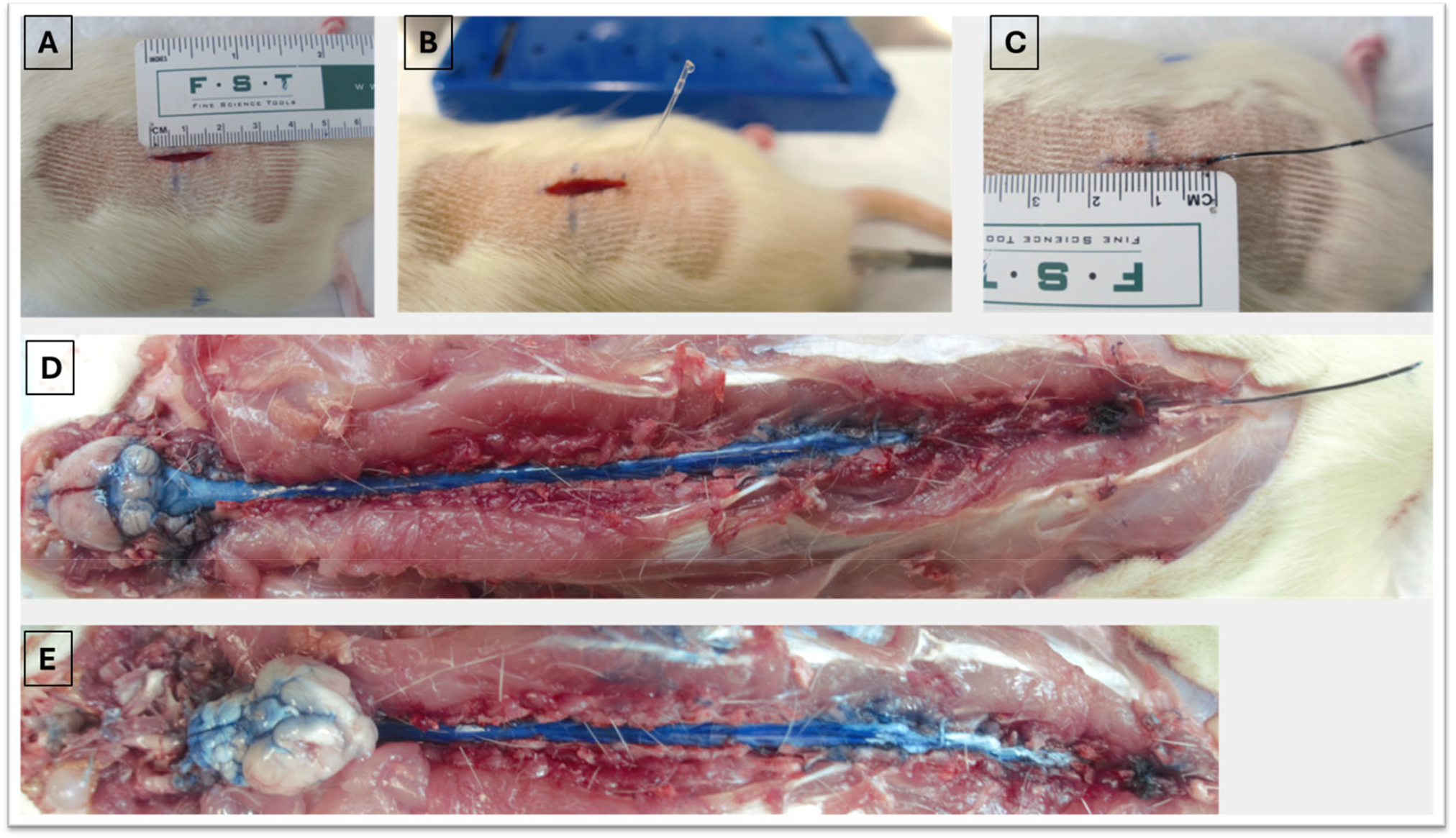
A: Mid-line skin incision (1.8 cm). B: After catheter placement, showing CSF drop on the catheter tip. C: Distal catheter connected to the needle tip while the dye is being injected. D: CNS dissection 90 minutes after injection, showing wide spread of the dye reaching the brain. E: The brain is inverted to show the dye in the brain basal CSF cisterns.

## Discussion

Treatment of central nervous system (CNS) disorders is complicated by the blood-brain barrier (BBB) and blood-spinal cord barrier, whose endothelial tight junctions and active efflux mechanisms restrict CNS penetration of diagnostics and therapeutics. The intrathecal (IT) route bypasses the BBB by delivering agents directly into the cerebrospinal fluid (CSF), enabling rapid CNS distribution, high local concentrations, and minimal systemic exposure [4]. In a preclinical setting, evaluating this route serves a dual translational purpose. For therapeutics explicitly intended for clinical intrathecal administration, it provides a faithful model of human delivery. Alternatively, for early-stage compounds ultimately intended for systemic administration but not yet optimized for BBB penetration, the route establishes an accessible screening window for proof-of-concept testing, justifying further investment in their development.

Chronic IT catheterization in rodents, first established by Yaksh and Rudy [5], has seen widespread adoption, though its application has been fraught with technical variability and significant morbidity. Modifications fall broadly into two categories: invasive surgical exposures (e.g., laminectomy) and “blind” percutaneous/interlaminar punctures [6, 12]. Laminectomy offers direct dural visualization but is low-throughput and induces tissue trauma, localized inflammation, and potential glial activation, confounding interpretation of neurological models (e.g., pain, SCI) [12]. Blind percutaneous methods (needle-through-needle, guide cannula) [9, 11] suffer from poor reliability and high rates of severe adverse events: direct injury to the spinal cord or cauda equina, subarachnoid hemorrhage from venous plexus disruption, and postoperative paresis or mortality [7, 12].

Our method overcomes these issues, integrating the precision of surgical exposure with the minimally invasive nature of an interlaminar puncture. The key innovation is microsurgical “under visual control” access to the L6-S1 interspace, which is anatomically distant from the conus medullaris and provides a wide margin of safety [11, 16]. Unlike blind methods relying on the tactile “pop” of the dura,[9] direct visualization lets the operator identify the dural midline precisely, avoid the epidural venous plexus, and prevent the hemorrhage that obscures the field or occludes the catheter.

We further identified pelvic padding to induce lumbar flexion, a common step, as a primary contributor to surgical failure: it produces a Valsalva-like effect, raising intra-abdominal pressure and congesting the epidural venous plexus. Our alternative, gentle rostral traction on the L6 spinous process, achieves sufficient flexion while preserving a clear, hemostatic field, making the “under visual control” approach feasible and reproducible. This precision is complemented by a smaller-bore (0.33 mm OD) polyurethane catheter, which reduces dural trauma and CSF leakage versus the larger PE-10 standard [9], and a muscle sheath closure that anchors the catheter without circumferential sutures that risk kinking or ischemic injury.

A persistent failure of lumbar IT delivery is unreliable rostral distribution: compounds pool in the lumbar cisternae due to complex CSF flow dynamics and fail to reach spinal and supraspinal targets [10, 17]. Our data address this directly. First, we implemented two controls for dose fidelity: dead volume compensation via “air-bubble tracking”, and epidural sealing with Surgifoam^®^ to prevent drug reflux (a major source of dose-response variability). Second, our Evans Blue dispersal data confirm that a bolus delivered at L6-S1 achieves full neuraxial distribution, including the ventral and dorsal brain cisterns, validating access to the global CSF circulation [8].

The most significant validation is pharmacodynamic target engagement in the brain. Reduced pS6/S6 ratios in the cerebral cortex after delivery of compound 8851 (an S6K1 inhibitor) show that our method can deliver a biologically active small molecule from the lumbar space to a distant supraspinal target at concentrations sufficient to modulate a specific pathway. Detecting brain phosphorylation changes as a proxy for kinase target engagement required refinement of traditional homogenization protocols: extraction and direct processing in hot 8% SDS buffer overcome the high lipid-to-protein ratio of neural tissue, which compromises protein resolution in brain samples. This prevents the sample aggregation and lane-smearing typical of brain Western blots, ensuring precise electrophoretic separation for accurate phosphoprotein quantification. The combined compound 8851 and Evans Blue results demonstrate the method’s generalizability and compatibility with diverse downstream analyses, addressing the recognized need for improved IT delivery methods [18].

This “under visual control” method requires a surgical microscope, foundational microsurgical skills, and higher instrument cost than percutaneous techniques. Nevertheless, the resultant reduction in animal morbidity and low procedural failure rate (10% in our hands, data not shown) outweigh the initial training investment. Future studies should validate this platform with other modalities such as protein- and nucleotide-based therapeutics, which have distinct CNS clearance and distribution properties [19].

## Conclusion

This study describes a safe, reliable, and reproducible protocol for lumbar IT catheterization that addresses key procedural and pharmacological challenges. Integration of precise anatomical targeting, controlled hemostasis, and dose calibration facilitates efficient CNS distribution. Refinements in tissue processing and Western blotting confirmed supraspinal pharmacodynamic target engagement by a small-molecule kinase inhibitor. Collectively, these methods can improve data quality, support animal welfare, and establish a robust platform for preclinical testing of CNS therapeutics.

## Supporting information

Online Resource 1

## Acknowledgments

We thank Dr. Jae Lee for his advice and guidance. This work was supported by the National Institutes of Health, The Buoniconti Fund, and The Miami Project to Cure Paralysis. V.P.L. holds the Walter G. Ross Distinguished Chair of Developmental Neuroscience.

## Author contributions

Conceptualization: O.S.E., J.L.B., V.P.L., H.A.; Investigation: O.S.E., V.P.L. B.A., A.B.M., H.A.; Methodology: O.S.E., J.L.B., V.P.L., H.A.; Writing – original draft: O.S.E., V.P.L. Writing – review and editing: O.S.E., A.B.M., B.A., J.L.B., V.P.L., H.A.

## Data availability

Data and metadata for the target engagement experiment is posted at the Open Data Commons – Spinal Cord Injury (https://odc-sci.org/) and will be made available upon acceptance of the paper.

## Ethics approval

All procedures, protocols, and medications were approved by the University of Miami Institutional Animal Care and Use Committee and were performed in accordance with the National Institutes of Health guide for the care and use of laboratory animals. Animal research at the University of Miami is accredited by the Association for Assessment and Accreditation of Laboratory Animal Care (AAALAC) International.

