## Supplementary material for "Lumbar intrathecal catheterization in rats targeting the cerebral cortex: a drug delivery method and validation": Online Resource 1

### Appendix 1

#### Surgical equipment and drugs

NON-WOVEN SPONGES (Dukal Corp.Ref. 7224)

Cotton-tip sticks

Artificial tears Ophthalmic ointment (PATTERSON Vet. NDC 14043-012-35)

WEBCRYL 4-0 surgical suture (Pivetal, cat. no. 21275161)

SURGIFOAM (ETHICON, cat. no. 4547284 Ref. 1977)

Sterile surgical blade no. 15 (Integra)

23-gauge × 1” needle

Isoflurane 250 ml (Isospire® by Dechra) anesthesia system equipped with an anesthesia induction chamber

Ketamine 100 mg/ml by (Ketamine Hydrochloride® by Dechra)

Xylazine 20 mg/ml. (Rompun® by Dechra)

Buprenorphine ER 1.3 mg/ml.

Buprenorphine HCL® 0.3 mg/ml

Heating pad controlled by a rectal thermometer.

Adson Forceps, 1 × 2 teeth (Fine Science Tools, cat. no. 11027-12)

Adson Forceps, Serrated (Fine Science Tools, cat. no. 11006-12)

Noyes Spring Scissors, 14-mm blades (Fine Science Tools, cat. no. 15012-12)

Olsen-Hegar Needle Holder (Fine Science Tools, cat. no. 12502-12)

Dumont no. 5/45 forceps, Dumoxel standard tip (Fine Science Tools, cat. no. 11251-35)

Scalpel handle no. 3 (Fine Science Tools, cat. no. 10003-12)

Electric shaver
